# Top-down corticostriatal gating of adaptive restraint during motivational conflict

**DOI:** 10.64898/2026.02.09.704889

**Authors:** Elizabeth Illescas-Huerta, Eduardo Hernández-Ortiz, Francisco Sotres-Bayon

**Author notes:** Correspondence (FS-B) and (EI-H).

## Abstract

Adaptive behavior under threat requires withholding reward pursuit when past experiences predict danger. Here, we identified a corticostriatal circuit that enables behavioral restraint during motivational conflict. Using a semi-naturalistic foraging task in male rats, we found that inactivation of the prelimbic cortex (PL) abolished restraint without impairing memory, and single-unit recordings revealed transient PL firing increases at decision points in conflict trials. c-Fos mapping showed selective engagement of the PL and nucleus accumbens core (NAcC) but not the shell. Inactivation further revealed that the anterior nucleus accumbens (aNAcC), but not posterior NAcC, is necessary for restraint, as posterior disruption broadly reduced reward-seeking regardless of threat. Silencing PL→aNAcC projections reproduced anterior NAcC effects without affecting memory or motivation, and this pathway was required under learned, but not innate, threats. These findings define a top-down corticostriatal mechanism in which prefrontal control of striatal activity specifically integrates learned threat and reward signals, enabling flexible decision-making and adaptive restraint in the face of danger.

## Introduction

Survival in dynamic environments requires urgent decisions that balance competing drives, such as the pursuit of essential rewards and the avoidance of potential threats^1,2^. In real-world scenarios, food and other resources often co-occur with danger-predictive cues, forcing animals to suppress their innate or learned defensive reactions to obtain vital outcomes. These approach–avoidance decisions are among the most complex that an organism can face, requiring animals to integrate past aversive experiences, anticipate future threats, and value rewards^3–6^. In humans, approach-avoidance decisions range from everyday risks to existential dilemmas, and are commonly impaired in conditions such as post-traumatic stress disorder, generalized anxiety, and depression, where individuals struggle to evaluate options in emotionally uncertain contexts^7–9^. Understanding the neural basis of these motivational conflicts is critical for elucidating how the brain guides adaptive behavior when a potential danger is anticipated. A key component of decision-making is behavioral restraint: the ability to withhold reward-seeking actions when doing so may incur a cost, such as a potential threat^10^. This internal restraining of behavior based on remembered danger lies at the core of adaptive responses under motivational conflict; however, the neural mechanisms supporting this capacity remain unclear.

In the context of motivational conflict, while the amygdala encodes threat-related information and orchestrates defensive behaviors^11–13^, striatal circuits, including the nucleus accumbens (NAc) and ventral pallidum (VP), integrate threat and reward signals to shape motivated actions^14–17^. In parallel, the medial prefrontal cortex (mPFC) in rodents contributes to action selection under conflicting motivational demands, partly through regulatory control of subcortical circuits. In particular, interactions between the mPFC and NAc have been implicated in approach–avoidance decision-making^18–23^. However, whether specific prefrontal-striatal projections mediate memory-guided restraint in response to learned threats remains unknown.

To address this gap, we employed the crossing-mediated conflict (CMC) task, an ecologically relevant foraging paradigm in which rats decide whether to cross a shock-paired grid to retrieve food, guided by auditory and visual cues signaling a reward or danger^17,24^. This paradigm models naturalistic decisions, in which reward pursuit must be weighed against potential risks, requiring animals to suppress defensive reactions to obtain rewards in a risky context. Using a multimodal approach, including pharmacological inactivation, exploratory single-unit recordings, c-Fos mapping, and optogenetic manipulation, we dissected the role of mPFC–NAc circuits in this process. We found that both the prelimbic (PL) region of the mPFC and the anterior NAc core (aNAcC) of the striatum were selectively recruited and necessary for behavioral restraint during conflict. In general, optogenetic silencing of the PL→aNAcC projection diminished this restraint, even in the presence of threat-predictive cues, without impairing memory or basic motivation. These findings reveal a corticostriatal mechanism that integrates conflicting motivational signals to guide adaptive actions under threat conditions.

## Results

Rats were trained in the CMC task to discriminate between safe (non-conflict) and risky (conflict) crossings to obtain food rewards. They learned to associate distinct cues with opposing outcomes, such as food or foot shock, and used this information to guide decisions in either safe or threatening contexts (Fig. 1A). The training protocol comprised three phases: (i) reward learning, with a light cue signaling food availability, (ii) threat learning, with a white noise cue paired with a foot shock, and (iii) discrimination training between non-conflict (light only) and conflict (light plus noise) trials. In the final test, no shocks were delivered, ensuring that the decisions reflected memory-guided integration of the learned reward and threat associations. The representative behaviors during this phase are shown in video 1.

**Fig. 1.**
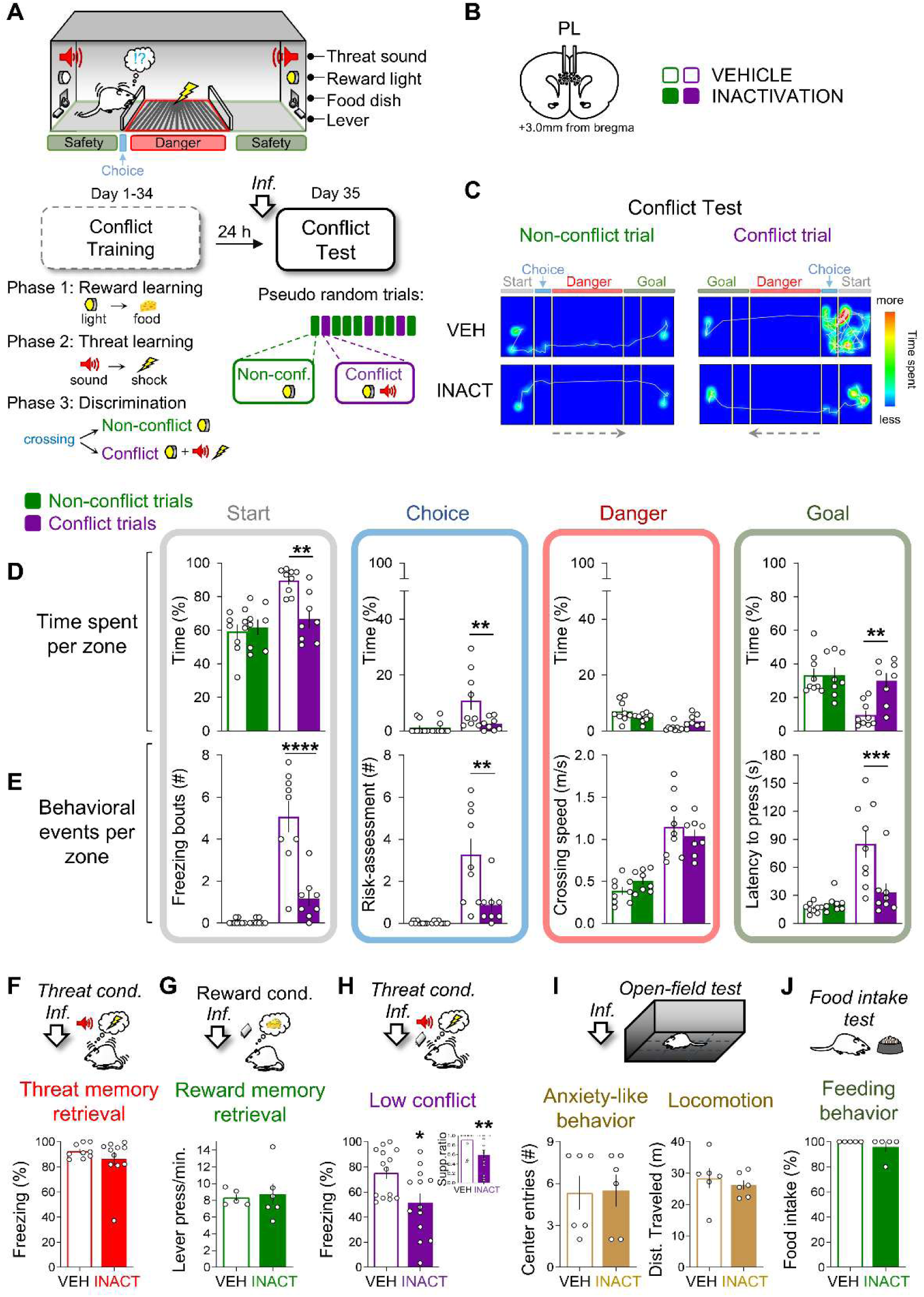
PL inactivation abolishes defensive responses and accelerates reward approach under learned threat. (**A, Top**) Schematic representation of the crossing-mediated conflict task. Rats foraged between feeding zones under non-conflict trials (reward cue only) and conflict trials (reward cue plus threat cue). (**A, Bottom**) Training comprised: (i) reward learning via light cue in a safe zone, (ii) threat learning via white noise in a danger zone, and (iii) discrimination between conflict (light + noise) and non-conflict (light only) trials. In the conflict test, rats decided whether to cross an electrified grid to obtain food. (**B**) Cannula placement in PL. Before testing, rats received PL infusions of saline (VEH, n = 9) or muscimol+baclofen (INACT, n = 8). (**C**) Representative movement tracking and heatmaps from VEH and INACT rats. (**D**) PL inactivation decreased the time spent in the start (F(1,15) = 4.66, P = 0.04; conflict P = 0.01) and choice zones (F(1,15) = 4.65, P = 0.04; conflict P = 0.009), and increased goal-zone time (F(1,15) = 5.46, P = 0.033, conflict P = 0.001) during conflict trials, with no effect in non-conflict trials. (**E**) In conflict trials, PL inactivation reduced freezing (F(1,15) = 20.17, P < 0.001; conflict, P < 0.0001) and risk assessment (F(1,15) = 7.42, P = 0.01; conflict, P = 0.001) in the start zone and decreased the latency to cross into the goal zone and press the lever (F(1,15) = 5.40, P = 0.034; conflict, P = 0.0008). (**F–G**) No effect on freezing (t(18) = 0.99, P = 0.331) or lever pressing (t(9) = 0.27, P = 0.79) in the isolated threat (VEH, n = 9; INACT, n = 11) or reward (VEH, n = 5; INACT, n = 6) retrieval tests. (**H**) Under low conflict, PL inactivation reduced freezing (t(25) = 2.68, P = 0.01) and the suppression ratio (t(25) = 2.97, P = 0.006) (VEH, n = 14; INACT, n = 13). (**I–J**) No differences in anxiety-like behavior, locomotion (t(10) = 0.10, P = 0.92; t(10) = 0.60, P = 0.56), or feeding (t(8) = 1.00, P = 0.34) (open field test, VEH, n = 6; INACT, n = 6; food intake test, VEH, n = 5; INACT, n = 5). Data are presented as the mean ± SEM. Empty circles represent individual rats. *P < 0.05, **P < 0.01, ***P < 0.001, ****P < 0.0001, Bonferroni post hoc after two-way mixed-design ANOVA or Student’s t-test.

### PL, but not IL, enables restraint under learned threat

The mPFC has been implicated in the regulation of defensive responses to threat-predictive cues, with the infralimbic (IL) cortex suppressing such responses and the PL facilitating them^25,26^. Based on this framework, we initially hypothesized that IL might be necessary to override defensive responses during reward-seeking under conflict. However, pharmacological inactivation of IL prior to a memory-guided conflict test did not alter behavior; rats spent similar time across spatial zones (start, choice, danger, goal) and showed unaffected measures of defensive behavior, including freezing, risk assessment, crossing speed, and lever-press latency (Fig. S1, A–D). These results suggest that IL is not required to restrain behavior when threat-related memories must be weighed against current goals.

In contrast, PL inactivation markedly impaired performance during the conflict trials (Fig. 1B-E). Rats spent significantly less time in the start zone and showed reduced freezing and risk assessment, along with shorter latencies to press the lever in the goal zone. Notably, the time spent and behavioral events in the danger zone were unaffected, indicating that PL inactivation did not cause generalized disinhibition or impair the ability to discriminate between zones. Instead, behavior during conflict trials became indistinguishable from that during non-conflict trials, suggesting that PL inactivation abolished behavioral restraint normally imposed by the learned threat. To rule out non-specific effects, we performed a series of control experiments using separate cohorts. PL inactivation did not alter freezing to a conditioned tone or instrumental food-seeking behavior (Fig. 1F-G), nor did it affect anxiety-like behavior, spontaneous locomotion, or food consumption (Fig. 1I-J). Furthermore, in a low-conflict version of the task, where approach costs were minimal because no threat-paired zones had to be crossed, PL inactivation reduced freezing and increased the suppression ratio (Fig.1H), suggesting that PL supports adaptive restraint even under low-demand conditions. These results demonstrated that PL activity, but not IL activity, is selectively required to suppress reward-seeking behavior when threat-predicting cues are present in the environment. Importantly, this effect cannot be explained by impairment in memory, motivation, or motor performance. Rather than directly mediating defensive responses, the PL appears to integrate competing motivational signals to guide behavior under conflict, consistent with prior reports implicating this region in cost–benefit evaluation, cognitive control, and flexible decision-making during approach–avoidance conflict^27–34^. Thus, PL supports context-sensitive restraint when memory-based threat signals must be overcome in the pursuit of rewards, setting the stage for downstream circuits that may implement cortical decisions.

### PL activity tracks decisions under threat and recruits a PL–NAcC circuit

Given that PL activity is required for behavioral restraint under conflict conditions, we next examined whether PL neurons were preferentially engaged under such conditions. We recorded extracellular single-unit activity in well-trained rats performing the conflict test (Fig. 2A-B), focusing on responses time-locked to the moment animals crossed from the choice zone to the danger zone. This analysis revealed distinct neuronal populations whose activity firing rate depended on the motivational context: a particular neuronal subpopulation responded selectively during conflict trials, others during non-conflict trials, and a smaller neuronal subset in both; with conflict-selective cells being more prevalent than expected by chance (Fig. 2C, top). Representative peri-stimulus time histograms (300 ms bins) show that conflict-responsive neurons often exhibit gradual ramping before zone entry, followed by a sharp peak at crossing and a rapid return toward the baseline. In contrast, non-conflict-responsive neurons typically showed more gradual modulations, frequently including peri-event inhibition, before a modest post-crossing increase (Fig. 2C, bottom panel). At the population level, conflict-responsive neurons displayed pronounced transient excitation centered on the crossing, typically preceded by ramping activity, whereas non-conflict-responsive neurons showed slower and less sharply tuned patterns (Fig. 2D). Quantification of the mean z-score activity (±600 ms) confirmed significantly higher firing in conflict neurons, and area-under-the-curve analyses revealed that most conflict-responsive neurons increased firing after the decision point, whereas non-conflict neurons were more often inhibited. This divergence was evident in both the proportion of excitatory and inhibitory responses (pie charts) and the direction of individual pre-/post-trajectories (color-coded lines) (Fig. 2E).

**Fig. 2.**
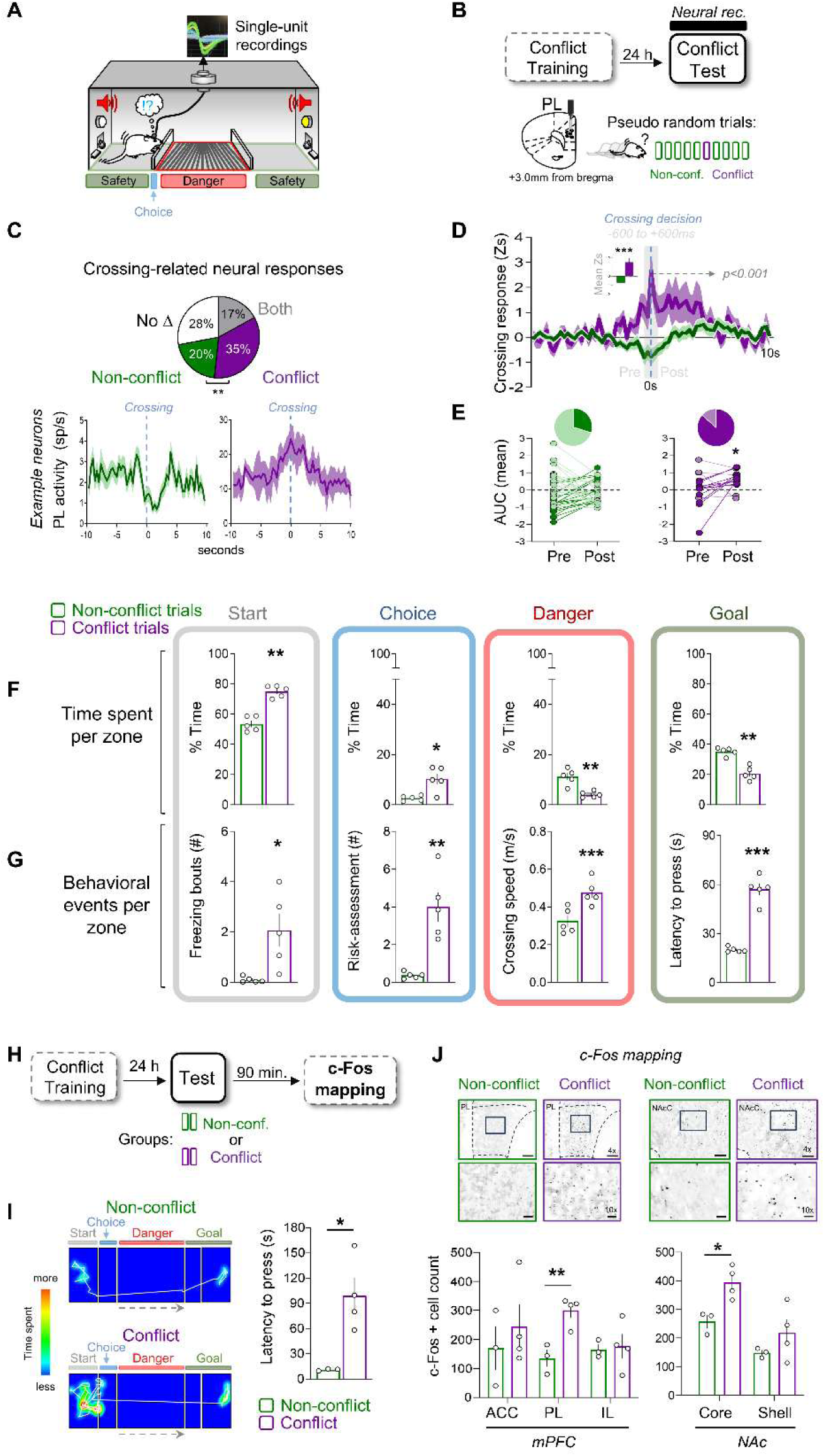
PL neuronal dynamics and c-Fos mapping identify selective recruitment of a PL-NacC circuit under threat. (**A**) Schematic of single-unit recordings in PL during the conflict test (n = 5 rats). (**B**) Electrode placement and design. Rats completed six trial blocks (nine non-conflict, one conflict). (**C, Top**) Proportions of PL neurons (n = 118) selectively responsive during conflict (n = 41), non-conflict (n = 24), both (n = 20), or unmodulated (n = 33) differed from chance (χ²(3) = 8.98, P = 0.0295). Conflict-responsive neurons were more prevalent than both dual-responsive (P = 0.0029) and non-conflict-responsive neurons (P = 0.0197). (**C, Bottom**) Representative neurons responsive in non-conflict (left) or conflict (right) trials. PSTHs (300 ms bins) aligned to crossing point (blue dashed line); shaded = ± SEM. (**D**) Average normalized firing rates (z-scores) of conflict- and non-conflict-responsive neurons. Conflict neurons (n = 15) showed ramping, a peri-entry peak, and rapid baseline return, whereas non-conflict neurons (n = 42) displayed slower, tonic patterns. Inset: mean activity (±600 ms) was higher in conflict than in non-conflict neurons (t(55) = 3.98, P = 0.0002). (**E**) Area under the curve confirmed excitation in conflict-excited neurons vs. baseline (t(14) = 2.16, P = 0.04), but no modulation was observed in non-conflict neurons (t(41) = 0.38, P = 0.69). Pie charts show excitation/inhibition in conflict (87%/13%) and non-conflict (29%/71%). (**F**) In conflict trials, rats spent more time in start (t(4) = 8.24, P = 0.001) and choice zones (t(4) = 2.92, P = 0.04), but less in danger (t(4) = 3.72, P = 0.02) and goal zones (t(4) = 6.83, P = 0.01) than in non-conflict trials. (**G**) Conflict trials increased freezing (t(4) = 2.29, P = 0.04), risk assessment (t(4) = 5.02, P = 0.007), speed (t(4) = 11.44, P = 0.0003), and the latency to cross into the goal zone and press the lever (t(4) = 10.22, P = 0.0005). (**H**) Rats (non-conflict, n = 4; conflict, n = 3) were perfused 90 min after the test for c-Fos mapping. (**I**) Representative trajectories and heatmaps; conflict rats had longer lever-press latencies (t(5) = 3.44, P = 0.01). (**J**) c-Fos mapping revealed higher PL (t(5) = 4.30, P = 0.007) and NAcC (t(5) = 3.58, P = 0.01) activity in conflict rats; no differences in ACC, IL, or NAcS. Representative PL and NAcC images shown. Data are mean ± SEM; *P < 0.05, **P < 0.01, ***P < 0.001, Student’s t-test unless noted.

These neuronal dynamics were accompanied by distinct behavioral patterns: during conflict trials, rats spent more time in the start and choice zones and less time in the danger and goal zones (Fig. 2F), and showed increased freezing, greater risk assessment, and longer lever-press latencies, despite rapid danger-zone crossings (Fig. 2G). Importantly, locomotor speed in the start (safe) zone did not differ between conditions (non-conflict: 0.029 ± 0.002 m/s; conflict: 0.022 ± 0.002 m/s; n = 5 rats; paired t-test, t(4) = 1.48, P = 0.21), arguing against generalized locomotor differences as an explanation for the decision-aligned PL activity. Given the known role of the NAc in mediating reward and aversion^35,36^, we assessed whether other brain regions were recruited during this form of behavioral restraint by quantifying c-Fos expression in rats exposed to either conflict or non-conflict trials (Fig. 2H). Consistent with the electrophysiological results, conflict-exposed rats displayed higher lever-press latencies (Fig. 2I) and increased c-Fos levels in the PL relative to controls, with no differences in the IL or anterior cingulate cortex (ACC). In the NAc, conflict exposure selectively increased c-Fos expression within the core (NAcC), but not in the shell (NAcS) (Fig. 2J). Together, these findings suggest that motivational conflict engages a distributed circuit linking the PL and NAcC, supporting their joint recruitment during memory-guided behavioral restraint. This corticostriatal interaction may underlie how top-down evaluations in the PL shape adaptive action selection in the face of threats.

### aNACc mediates restraint under learned threat

Our c-Fos mapping revealed selective activation of the NAcC, but not the NAcS, during behavioral restraint under conflict, leading us to functionally target the NAcC. Pharmacological inactivation of the entire NAcC prior to the conflict test produced divergent behavioral outcomes (Fig. S2, A). Some rats showed reduced latencies to press under conflict, while others displayed generalized suppression of approach behavior across trials (Fig. S2, B). Based on previous reports of functional heterogeneity along the anteroposterior axis of the accumbens^37^, we applied an unsupervised clustering approach, using cannula placement in the NAcC. This cluster analysis identified two subgroups corresponding to anterior (+2.5 to +2.0 mm) and posterior (+2.0 to +1.6 mm) cannula placement in the NAc (Fig. S2, C). In contrast, PL inactivation resulted in a uniform and clustered behavioral profile. These findings prompted us to analyze the anterior and posterior NAcC separately in the following experiments, beginning with the anterior NAcC (aNAcC), which is the region associated with reduced restraint in cluster analysis.

To determine the specific contribution of aNAcC, we pharmacologically inactivated this region before the conflict test. aNAcC inactivation did not affect the time spent in any of the spatial zones (Fig. 3A–C). However, during conflict trials, it significantly decreased freezing and risk assessment, resulting in shorter lever-press latencies (Fig. 3D). These changes specifically reduced freezing and risk assessment, and shorter lever press latencies, resembled the effects of PL inactivation, although they were somewhat milder, and occurred without impairing spatial navigation or trial discrimination, as indicated by the preserved crossing speeds. These results suggest that aNAcC activity helps suppress premature reward pursuit when a threat memory is present. To further assess specificity, we tested aNAcC-inactivated rats using control tasks. Freezing to a conditioned tone and instrumental food-seeking behavior were unaffected (Fig. 3E–F), indicating that basic threat and reward memories remained intact. However, in a low-conflict retrieval task in which food-seeking occurs without crossing, but in the presence of a threat-predictive cue, aNAcC inactivation reduced freezing and increased the suppression ratio (Fig. 3G), supporting its role in modulating behavior, even when motor demands are minimal. Anxiety-like behavior, locomotion, and feeding also remained unchanged (Fig. 3H–I). Together, these findings identify aNAcC as a critical node for implementing behavioral restraint during motivational conflict. Its contribution parallels that of the PL, supporting the idea that prefrontal and striatal regions interact to regulate adaptive decision-making when threat and reward information must be integrated.

**Fig. 3.**
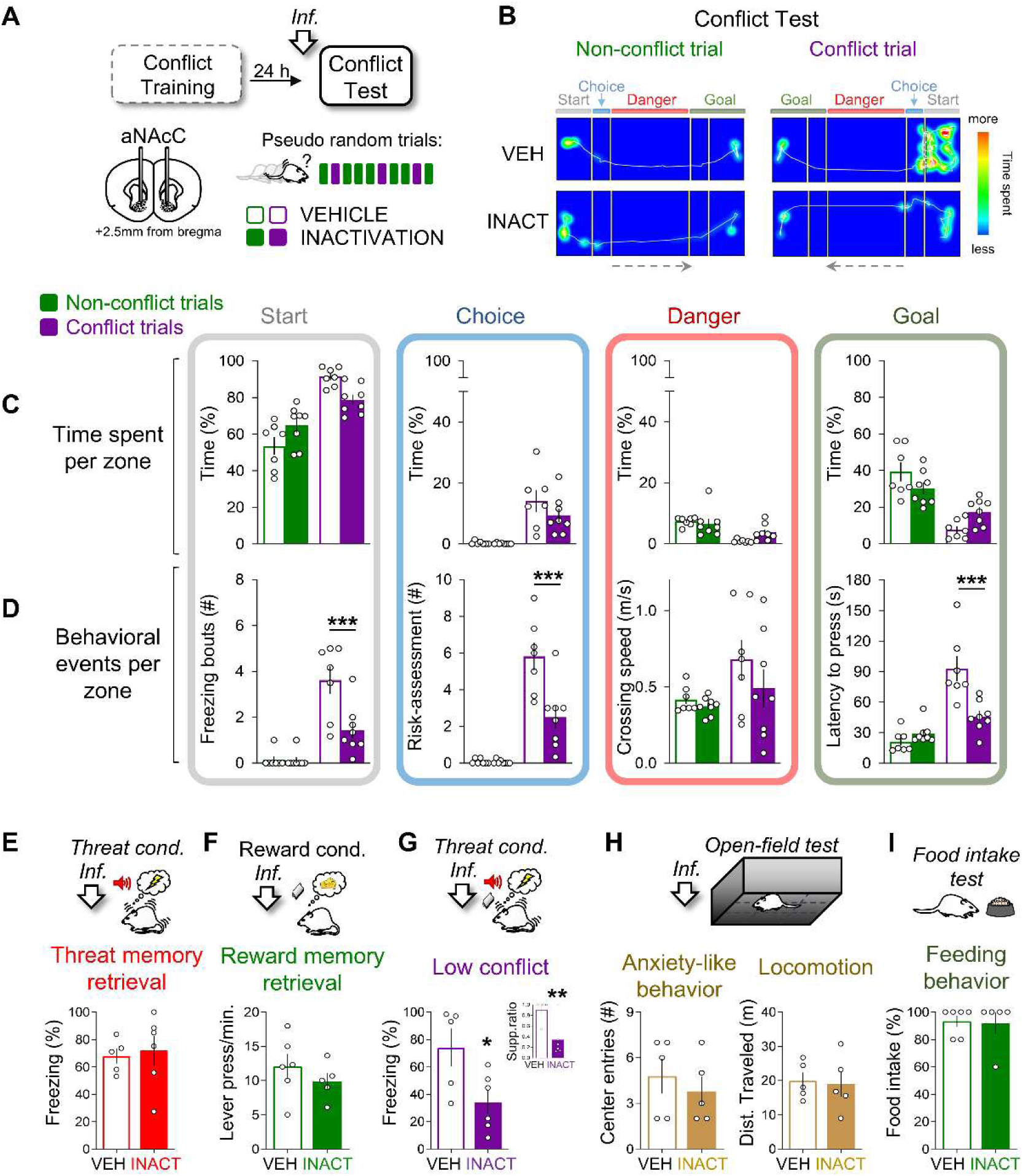
aNAcC inactivation reduces defensive responses without affecting threat or reward memory. (**A**) Cannula placements in anterior NAcC (aNAcC). Rats received vehicle (VEH, n = 7) or muscimol+baclofen (INACT, n = 8) before the conflict test. (**B**) Representative movement tracking and heatmaps from VEH and INACT rats during the first two consecutive non-conflict and conflict trials. (**C**) aNAcC inactivation did not produce a main group effect on the time spent in the start (F(1,13) = 0.02, P = 0.87), choice (F(1,13) = 1.54, P = 0.23), danger (F(1,13) = 0.63, P = 0.44), or goal zones (F(1,13) = 0.008, P = 0.92) despite a significant interaction effect. (**D**) In conflict trials, inactivation reduced freezing (F(1,13) = 9.94, P = 0.007; conflict P = 0.0004) and risk assessment (F(1,13) = 11.83, P = 0.004; conflict P = 0.0001) in the start zone and decreased the latency to cross into the goal zone and press the lever (F(1,13) = 5.57, P = 0.03; conflict P = 0.0001). (**E–F**) No effect on freezing (t(9) = 0.30, P = 0.76) or lever pressing (t(9) = 0.94, P = 0.36) in the isolated threat (VEH, n = 5; INACT, n = 6) or reward (VEH, n = 6; INACT, n = 5) retrieval tests. (**G**) Under low conflict, inactivation decreased freezing (t(9) = 2.53, P = 0.03) and the suppression ratio (t(9) = 3.30, P = 0.009) (VEH, n = 5; INACT, n = 6). (**H–I**) No differences in anxiety-like behavior or locomotion in the open field (VEH, n = 5; INACT, n = 5) or feeding in the free-food intake test (VEH, n = 6; INACT, n = 5). Data are presented as mean ± SEM; *P < 0.05, **P < 0.01, ***P < 0.001, Bonferroni post-hoc test after two-way mixed-design ANOVA or Student’s t-test.

### pNAc promotes reward-seeking regardless of threat

Next, we examined the role of the posterior nucleus accumbens core (pNAcC), identified by cluster analysis as associated with enhanced behavioral restraint. Pharmacological inactivation of this region prior to the conflict test (Fig. 4A) did not alter time allocation across spatial zones, defensive responses (freezing, risk assessment), or crossing speed during conflict or non-conflict trials (Fig. 4B–D). However, it significantly increased latency to press the lever for food, regardless of trial type. Notably, 40% of the pNAcC-inactivated rats failed to cross the arena within 180 s, indicating a reduction in baseline incentive motivation rather than a trial-type-specific effect. This motivational deficit persisted even under simplified conditions, in which rats were placed directly in the goal zone without the need to cross. In this context, pNAcC-inactivated animals were slower to press the lever (VEH: 4s, INACT: 54s; t(14) = 2.76, P = 0.01), similar to the reduced reward pursuit observed with VP inactivation^17^. This suggests that pNAcC and VP may work in concert to sustain an appetitive drive independent of the threat context.

**Fig. 4.**
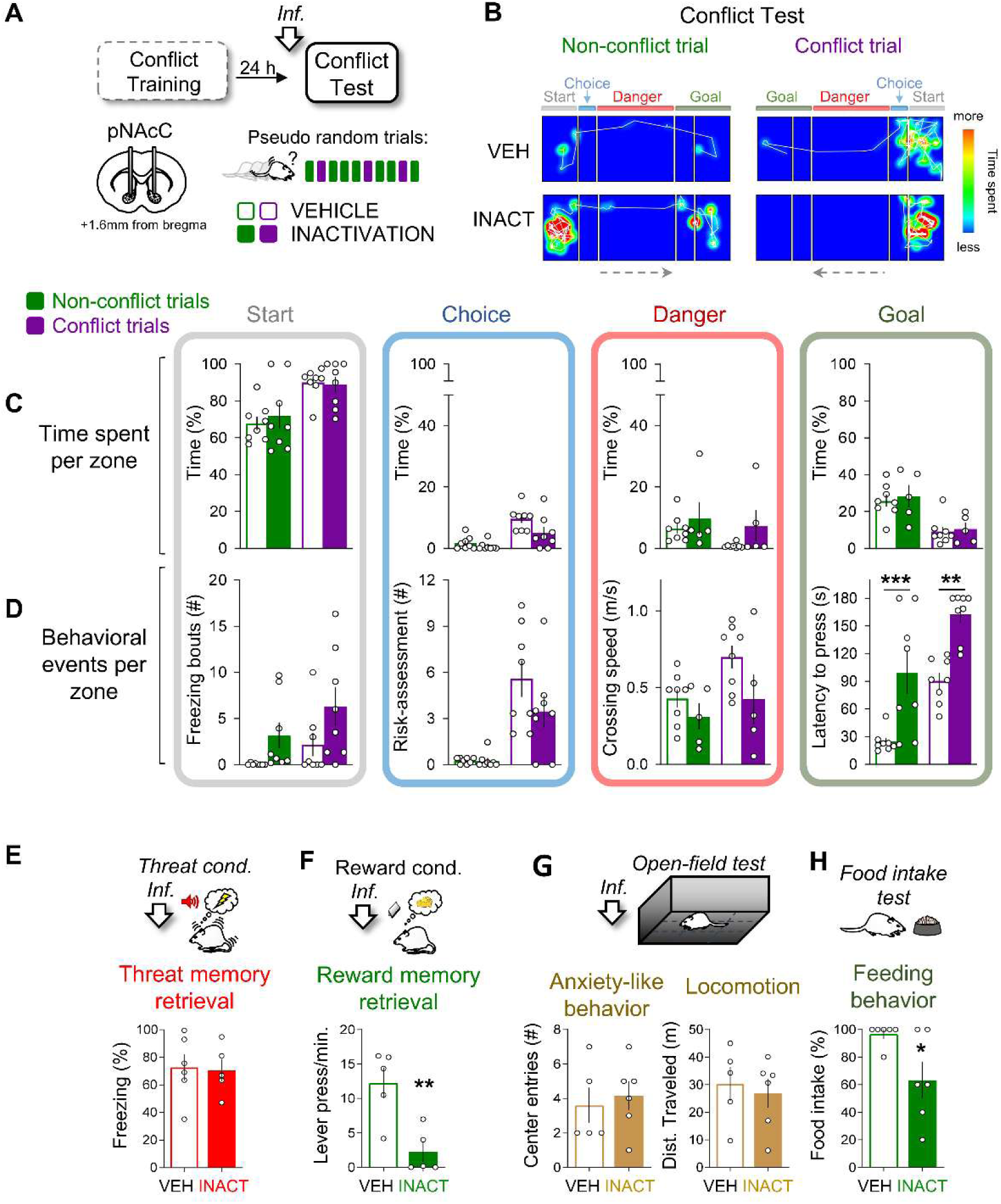
pNAcC inactivation impairs general reward pursuit regardless of threat signals. (**A**) Cannula placements in posterior NAcC (pNAcC). Rats received vehicle (VEH, n = 8) or muscimol+baclofen (INACT, n = 8) before the conflict test. (**B**) Representative movement tracking and heatmaps from VEH and INACT rats during non-conflict (reward cue only) and conflict (reward cue + threat cue) trials. (**C**) pNAcC inactivation did not significantly alter time in start (F(1,14) = 0.06, P = 0.80), choice (F(1,14) = 4.24, P = 0.06), danger (F(1,11) = 1.45, P = 0.25), or goal zones (F(1,11) = 0.19, P = 0.67). (**D**) Inactivation increased the latency to cross into the goal zone and press the lever (F(1,14) = 25.18, P = 0.0002; planned comparison conflict, P = 0.0013; non-conflict, P = 0.0009). (**E**) No effect on freezing in isolated threat memory retrieval (t(9) = 0.15, P = 0.87; VEH, n = 6; INACT, n = 5). (**F**) In reward memory retrieval, inactivation reduced lever pressing (t(8) = 3.75, P = 0.005; VEH, n = 5; INACT, n = 5). (**G**) No differences in anxiety-like behavior (t(9) = 4.33, P = 0.67) or locomotion (t(9) = 0.41, P = 0.68) were observed in the open field (VEH, n = 5; INACT, n = 6). (**H**) Inactivation reduced feeding in the free food intake test (t(10) = 2.46, P = 0.03; VEH, n = 6; INACT, n = 6). Data are presented as mean ± SEM; *P < 0.05, **P < 0.01, ***P < 0.001, Bonferroni post-hoc test after two-way mixed-design ANOVA or Student’s t-test.

To further confirm that the observed behavioral changes reflected motivational rather than mnemonic deficits, we conducted additional tests. For instance, pNAcC inactivation did not impair freezing responses to the conditioned tone (Fig. 4E), indicating an intact threat memory. However, reward-related behaviors were consistently reduced: pNAcC-inactivated rats pressed less for food (Fig. 4F) and consumed less food in the home cage (Fig. 4H), similar to what was observed in the caudal NAcS in a previous report^37^. The measures of anxiety-like behavior and locomotion remained unchanged (Fig. 4G). Together, these results indicate that the pNAcC is critical for sustaining goal-directed reward seeking, independent of threat or behavioral restraint, underscoring the functional dissociation along the rostrocaudal axis of the NAcC.

### PL→aNAcC pathway enables adaptive restraint under learned threat

Our pharmacological inactivation findings demonstrated that both PL and aNAcC are necessary for suppressing food-seeking behavior in the presence of learned threat cues, suggesting that a prefrontal–striatal circuit supports behavioral restraint. To directly test the role of the PL→aNAcC projection, we used optogenetic tools to selectively inhibit or activate this pathway during the conflict task. We bilaterally infused PL with an adeno-associated virus (AAV) encoding both archaerhodopsin (Arch) and enhanced yellow fluorescent protein (eYFP) under the control of the calcium calmodulin Kinase II (CaMKII) promoter to target glutamatergic neurons. To validate the photoinhibition efficacy, we implanted a unilateral optrode into the aNAcC of rats. Photoinhibition of PL terminals in aNAcC significantly reduced the firing rate of over one-third of the recorded neurons, confirming the effective suppression of PL input to aNAcC (Fig. 5A). Anatomical verification confirmed robust eYFP expression in PL pyramidal neurons and dense terminal labeling in the aNAcC, consistent with targeted pathway expression (Fig. 5B).

**Fig. 5.**
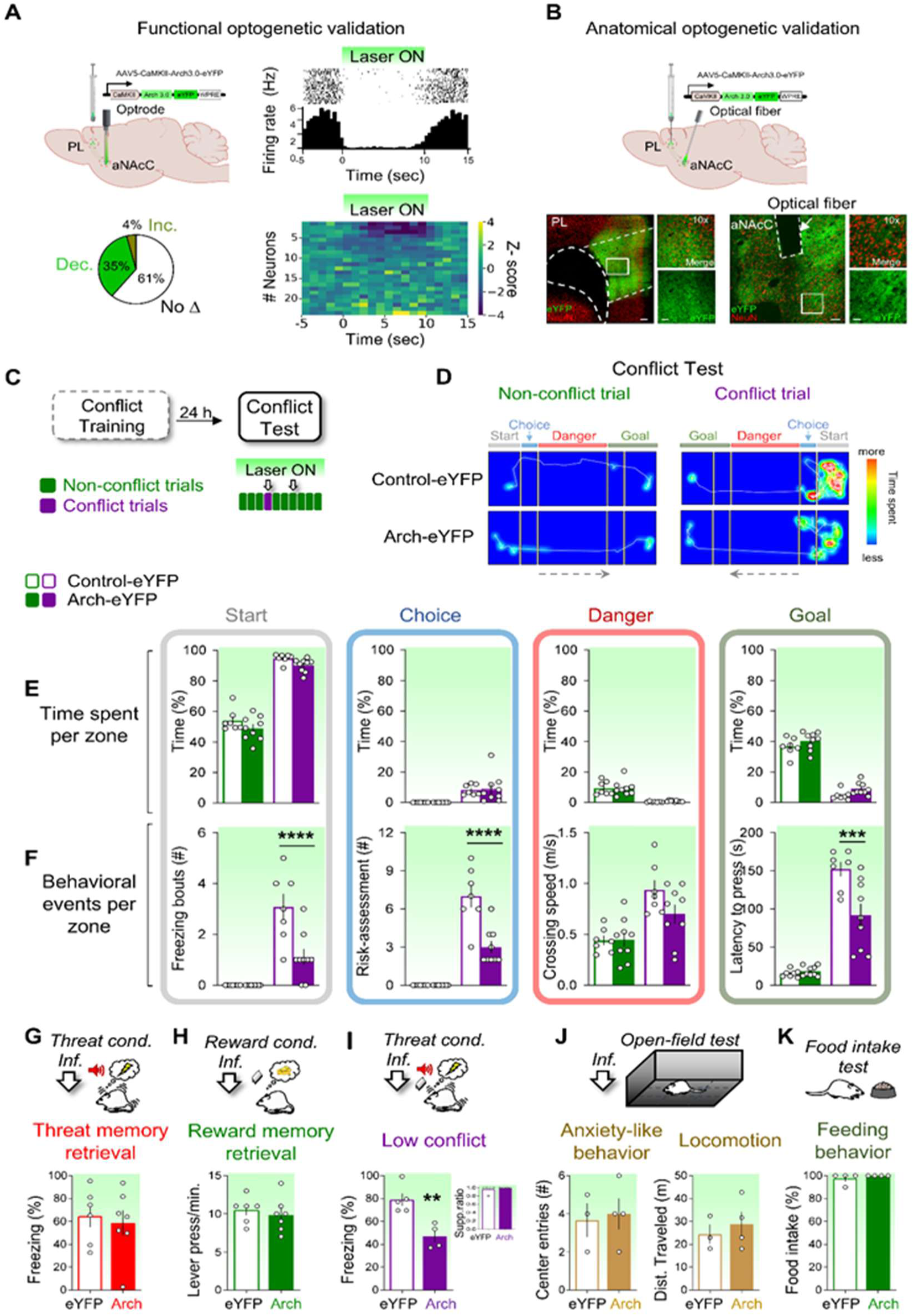
Silencing PL→aNAcC projection reduces defensive responses and facilitates reward approach. (**A, Top left**) Schematic showing viral injection of AAV5-CaMKII-Arch-EYFP in the PL and optrode implantation in the aNAcC (n = 5 rats). (**A, Top right**) Raster plot of a representative unit during photoinhibition (laser on). (**A, Bottom left**) Percentage of PL→aNAcC neurons (n = 23) modulated during photoinhibition. The firing rate decreased in eight (35%) neurons and increased in one (4%) (Wilcoxon signed-rank test, 1 s bins, all P < 0.05). **(A, Bottom right)**, Color map (z-score) of neural responses aligned to laser onset. (**B, Top**) Schematic representation of viral infection in the PL and bilateral optical fiber implantation in the aNAcC. (**B, Bottom**) Micrographs showing viral expression in the PL and axonal projections to the aNAcC (arrow = fiber track). (**C**) During the conflict test (Control-eYFP n = 7, Arch-eYFP n = 9), rats underwent two crossing trial blocks: laser ON followed by laser OFF (see Fig. S3). (**D**) Overlaid movement tracking and heatmaps from representative Control-eYFP and Arch-eYFP rats during the first two non-conflict and conflict trials. (**E**) Photoinhibition of PL→aNAcC projections did not alter zone times during the conflict test. (**F**) Photoinhibition reduced freezing (F(1,14) = 11.70, P = 0.004; conflict P < 0.0001) and risk assessment (F(1,14) = 18.38, P < 0.001; conflict P < 0.0001) in the start zone and decreased the latency to cross into the goal zone and press the lever (F(1,14) = 7.49, P = 0.01; conflict P = 0.0003) without affecting non-conflict trials. (**G–H**) No effect on freezing (t(11) = 0.39, P = 0.69) or lever pressing (t(11) = 0.56, P = 0.58) during the threat (Control-eYFP, n = 6; Arch-eYFP, n = 7) or reward (Control-eYFP, n = 6; Arch-eYFP, n = 7) retrieval tests. (**I**) In the low-conflict test, photoinhibition decreased freezing (t(7) = 4.13, P = 0.004) without altering the suppression ratio (t(7) = 0.88, P = 0.40; Control-eYFP, n = 5; Arch-eYFP, n = 4). (**J–K**) No differences in anxiety-like behavior (t(5) = 0.27, P = 0.79), locomotion (t(5) = 0.59, P = 0.57), or feeding (t(6) = 1.00, P = 0.35) were observed in the open field (Control-eYFP, n = 3; Arch-eYFP, n = 4) or free food intake tests (Control-eYFP, n = 4; Arch-eYFP, n = 4). Scale bars: 500 μm (PL) and 100 μm (NAcC); insets: 50 μm (PL) and 20 μm (NAcC). Data are presented as mean ± SEM; **P < 0.01, ***P < 0.001, ****P < 0.0001, Bonferroni post-hoc test after two-way mixed-design ANOVA or Student’s t-test.

For behavioral testing, we expressed either Arch or eYFP in PL neuronal bodies and bilaterally implanted optical fibers into the aNAcC of rats. Rats were tested in two consecutive trial blocks: one conflict and one non-conflict trial with photoinhibition (laser-on; Fig. 5C), followed by the same trial types without stimulation (laser-off; Fig. S3,A). PL→aNAcC photoinhibition selectively reduced defensive responses (freezing in the start zone and risk assessment in the choice zone) during conflict trials, with no effect on non-conflict behavior. These changes were accompanied by shorter latencies in pressing the goal zone. Trial discrimination and navigation remained intact, as indicated by the preserved crossing speed and spatial zone occupancy (Fig 5E-F). In laser-off trials, Arch-expressing rats exhibited longer latencies than eYFP-controls, possibly reflecting the delayed extinction of threat-associated memory (Fig. S3,B-C). To rule out general deficits in memory or motivation, threat, and reward retrieval were assessed using separate tasks. PL→aNAcC photoinhibition did not alter freezing to a conditioned tone or levers pressing for food (Fig. 5G-H), indicating intact cue-reward and cue-threat associations. In a low-conflict retrieval test, photoinhibition reduced freezing during laser-on tone presentation but not during the subsequent laser-off tone (Fig. S3,F), confirming a transient and specific effect. However, the suppression ratio remained unaltered during laser-on tone presentation, suggesting that photoinhibition was insufficient to elicit reward-seeking behavior, despite the suppression of passive defensive reactions in a low conflict (Fig. 5I). Anxiety-like behavior, locomotion, and feeding were unaffected (Fig. 5J–K).

To assess whether PL→aNAcC activity is sufficient to influence restraint, we expressed channel rhodopsin II (ChR2) or eYFP in PL neuronal soma and optically stimulated aNAcC terminals during the conflict task (Fig. S4, A). Contrary to the induction of selective restraint, photoactivation disrupted behavior more broadly; it increased crossing latency in both conflict and non-conflict trials without altering defensive behaviors (Fig. S4, C–F). Despite using the stimulation parameters used in a previous report^23^, the trajectory data revealed disorganized time allocation across zones (Fig. S4, B), suggesting that activation of the pathway impairs goal-directed behavior, regardless of the threat context. Together, these findings indicate that the PL→aNAcC projection is necessary for expressing adaptive behavioral restraint under threat, but that overactivation disrupts goal pursuit, highlighting the need for balanced activity in this corticostriatal pathway for decision-making under motivational conflict.

### aNAcC mediates restraint under innate threat independently of PL

Our previous results demonstrated that PL→aNAcC projection is necessary to restrain reward-seeking behavior in the presence of learned threat cues. However, in natural environments, animals must also be regulated in the presence of innate threats, such as bright and exposed spaces that signal danger during foraging. To examine whether the same prefrontal-striatal circuits support restraint under innate threat, we tested the effects of PL-aNACC inactivation as well as PL→aNAcC photoinhibition using the foraging-mediated innate conflict task^24^.

This task presents food at the center of a brightly lit open field, confronting the foraging drive with a natural aversive stimulus (Fig. 6A). The arena was divided into three zones: safe (periphery), dangerous (center), and choice (transition zone between them). Animals were tested in either conflict trials (food present) or non-conflict trials (no food present), with different groups used for each condition. Pharmacological inactivation of PL or aNAcC was performed before testing. Representative movement traces and heatmaps illustrate the zone occupancy during both trial types (Fig. 6B). PL inactivation did not alter the time spent in any zone during conflict or non-conflict trials, suggesting that PL activity was not required to restrain foraging in response to innate threats (Fig. 6C). This result aligns with previous findings that PL inactivation does not modulate innate defensive responses such as those evoked by predator exposure^38^. In contrast, aNAcC inactivation significantly reduced the time spent by rats in the safe zone and increased the time spent in the danger zone during conflict trials, with no effect on non-conflict trials (Fig. 6D). These results indicate that the aNAcC is necessary to constrain reward-seeking behavior when innate threat cues are present. The behavioral effects closely resemble those observed following systemic diazepam administration in the same task^24^, suggesting that the aNAcC integrates conflicting motivational signals across different threat domains in the same way.

**Fig. 6.**
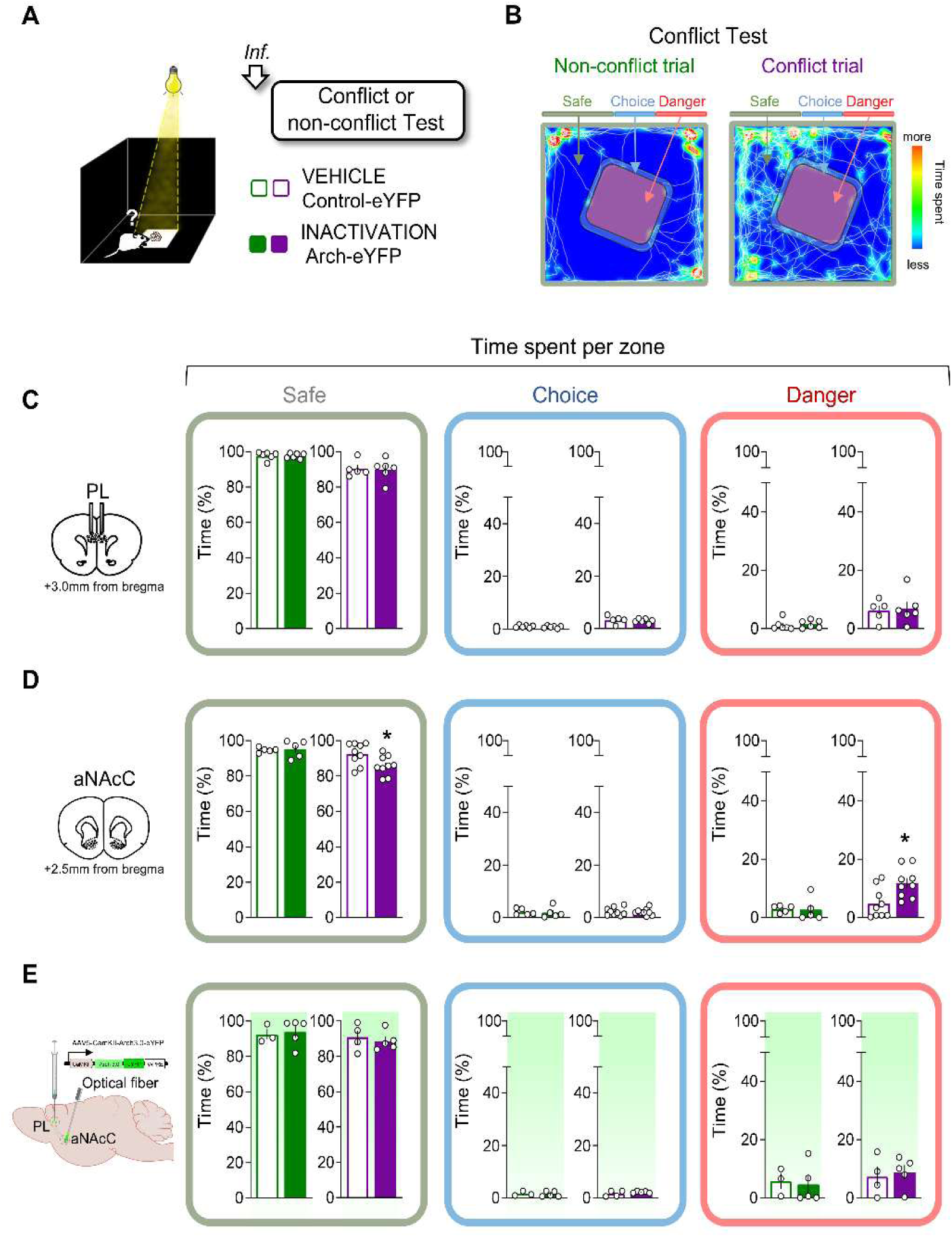
aNAcC, but not PL or PL→aNAcC projections, supports restraint under innate threat. (**A**) Rats were tested in a dark arena where they foraged for food in a brightly lit center (conflict test) versus an unlit center (non-conflict test). Before testing, they received vehicle (VEH) or muscimol+baclofen (INACT) infusions into the PL or aNAcC. (**B**) Overlaid movement tracking and heatmaps from a representative saline-infused rat in non-conflict and conflict trials. (**C**) Injection sites in the PL. Inactivation did not affect time spent in start (non-conflict: t(10) = 0.14, P = 0.88; conflict: t(9) = 0.11, P = 0.91), choice (non-conflict: t(10) = 0.79, P = 0.44; conflict: t(9) = 0.54, P = 0.59), or danger zones (non-conflict: t(10) = 0.38, P = 0.70; conflict: t(9) = 0.26, P = 0.79) (VEH, n = 6; INACT, n = 6 in non-conflict; VEH, n = 5; INACT, n = 6 in conflict). (**D**) Injection sites in the aNAcC. Inactivation reduced time in the start zone (t(16) = 2.42, P = 0.02) and increased time in the danger zone (t(16) = 2.83, P = 0.01) during conflict (VEH, n = 9; INACT, n = 9), with no effects in non-conflict (start: t(8) = 0.24, P = 0.81; danger: t(8) = 0.15, P = 0.88, VEH, n = 5; INACT, n = 5). (**E**) Viral injection in the PL and bilateral optical fiber implantation in the aNAcC. Photoinhibition of PL→aNAcC projections did not alter zone occupancy in non-conflict (Control-eYFP, n = 3; Arch-eYFP, n = 5) or conflict (Control-eYFP, n = 4; Arch-eYFP, n = 5) trials. Data are presented as mean ± SEM; *P < 0.05, Student’s t-test. The empty circles represent individual rats.

Given that previous studies have implicated prefrontal-striatal circuits in innate-avoidance behaviors^39,40^, we evaluated whether the PL→aNAcC projection is also involved in innate-threat restraint by expressing Arch or eYFP in the PL and bilaterally implanting optical fibers in the aNAcC (Fig. 6E). During the foraging conflict task, photoinhibition of the PL→aNAcC projection had no effect on zone occupancy under either conflict or non-conflict conditions. This suggests that, while aNAcC contributes to innate threat modulation, its regulation may involve inputs from brain regions other than the PL cortex. In summary, our findings indicate that the PL→aNAcC pathway is selectively engaged in learned-threat contexts to mediate behavioral restraint, whereas innate-threat regulation relies on distinct afferents to aNAcC. These results highlight the flexible recruitment of neural circuits for adaptive behavioral control depending on the nature of the threat to the individual.

## Discussion

Our results identified a distributed yet organized corticostriatal circuit that supports adaptive restraint under threat. The PL and aNAcC converge to withhold reward-seeking in the presence of learned threat cues, integrating competing motivational drives. In contrast, pNAcC regulates general incentive motivation regardless of threat. Notably, aNAcC also mediates restraint under innate threats, independent of PL input, indicating a shift in circuit recruitment depending on the threat source. These findings define a modular and flexible corticostriatal architecture for regulating decision-making under threat, with circuit engagement shaped by both threat type and task demands (Fig. 7). Together, these pathways form a top-down corticostriatal mechanism that supports adaptive reward pursuit under threat conditions.

**Fig. 7.**
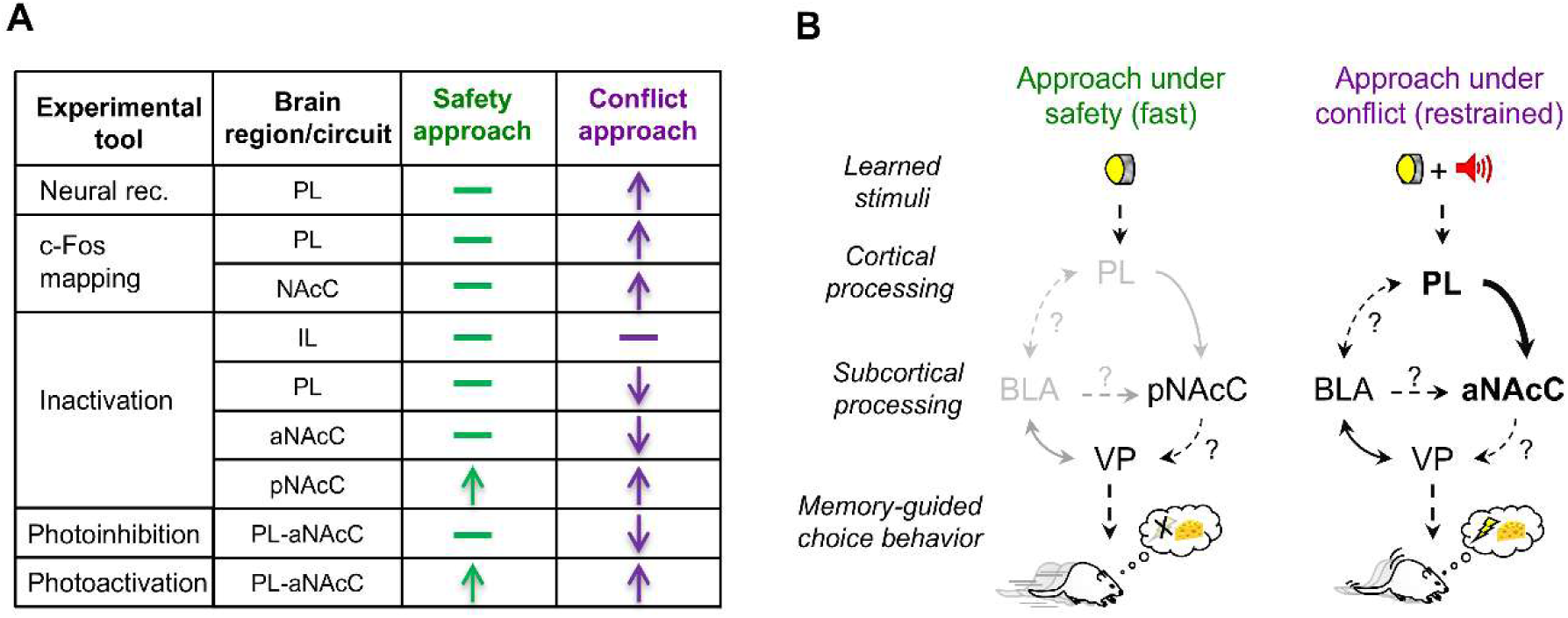
Experimental tools and proposed corticostriatal circuits for reward approach under safety vs. conflict. (**A**) Summary table of experimental tools (neural recordings, c-Fos mapping, inactivation, photoinhibition, and photoactivation) and the brain regions tested for each condition: safety approach (green) vs. conflict approach (purple). (**B**) Proposed circuit model. Left: In safe conditions, reward-predictive cues elicit fast, unrestrained approach behavior, primarily driven by motivational signals through the posterior nucleus accumbens core (pNAcC; this study) and ventral pallidum (VP¹⁷). Right: When reward and threat cues co-occur, animals display a restrained approach marked by hesitation and risk assessment. This adaptive behavior depends on the prelimbic cortex (PL) and its projection to the anterior NAc core (aNAcC; this study), likely with modulatory inputs from the basolateral amygdala (BLA) to the VP¹⁷. Together, these pathways form a top-down corticostriatal mechanism that supports adaptive reward pursuit under threat.

By combining pharmacological, electrophysiological, immunohistochemical, and optogenetic approaches in a semi-naturalistic foraging task, we found that the PL and aNAcC are co-engaged when rats decide whether to cross a risky zone for food. This ethologically grounded design, which allowed us to model decisions integrating memory, motivation, and internal conflict, captured components difficult to study in simpler organisms or constrained paradigms. While prior studies have implicated the prefrontal cortex and NAc in reward processing^41–45^ and aversive learning^30,46–50^, our results uniquely demonstrate the dynamic interplay between the PL and aNAcC, specifically during conflict involving a learned threat. Heightened activity in these regions during conflict was required to preserve the hesitation or restraint typically seen in response to remembered danger, without impairing motor output, motivation, or memory. This aligns with evidence that prefrontal networks can represent emotionally significant associations to guide flexible behavior^34^, and our findings identify a specific prefrontal–striatal pathway by which prior aversive learning exerts internal control over goal pursuit even in the absence of immediate danger, paralleling human findings on prefrontal–amygdala regulation of fear and reward^51,52^.

Within this pathway, PL emerges as a key hub in adaptive restraint. In conflict conditions, both single-unit and c-Fos measurements demonstrated enhanced PL activity relative to non-conflict trials. Earlier research suggested a generalized role of the PL in fear expression^26,27,38,42,47,53–55^; our data point to a more specific function: modulating approach behavior when learned threats must be overcome to obtain a reward. PL inactivation selectively disrupted hesitation during conflict without affecting non-conflict performance or simpler Pavlovian and instrumental behaviors. Recordings revealed temporally precise, conflict-specific activity—often ramping before zone entry, peaking sharply at the decision point, and returning rapidly to baseline— that was overrepresented relative to chance. Together with comparable start-zone speed across conditions, these temporally locked dynamics are most consistent with a context-dependent action-selection signal rather than generalized locomotor drive. These patterns, together with c-Fos data, argue against a role in general arousal or motor control, instead linking PL activity to a discrete signal for behavioral restraint. PL inactivation abolished defensive responses and sped up approach, complementing prior reports on approach–avoidance regulation^30,32^ and supporting a context-dependent role engaged during cue-driven but not innate threat^56^. NAcC manipulation revealed functional specialization along the anteroposterior axis. Inactivation of aNAcC reduced defensive responses and accelerated the approach during conflict, whereas pNAcC inactivation reduced reward pursuit, even in safe trials. This dissociation challenges the view of the NAcC as functionally homogeneous and suggests that the aNAcC acts as a context-sensitive gatekeeper for restraint, whereas the pNAcC sustains goal-directed behavior independent of threat^37,57^. Unlike PL, aNAcC participates in restraining both learned and innate threats, indicating its broader and more context-independent role.

Projection-specific optogenetics showed that inhibiting PL→aNAcC during conflict reduced defensive responses and shortened decision latency, directly confirming its causal role and mirroring the effects of aNAcC inactivation while sparing innate threat responses. This specificity underscores the precise pathway for cue-driven restraint and highlights potential targets for treating disorders characterized by impaired inhibition^58^. Conversely, activating this pathway disrupted goal-directed behavior even in non-conflict trials, suggesting that while PL→aNAcC pathway is necessary for adaptive restraint, excessive drive impairs action selection and underscores the importance of balanced corticostriatal signaling. These results may have translational implications for clinical practice. Pursuing rewards despite remembered threats—a behavior often described as “courage” in ethological and clinical contexts,^51,52^—likely engages overlapping prefrontal–amygdala–striatal networks in humans. Mapping this circuitry provides a framework for studying approach–avoidance conflict across species, and for modeling maladaptive decisions, including addiction, where the task could incorporate drug rewards to investigate relapse mechanisms.

While our study establishes the necessity of the PL→aNAcC pathway, further work is needed to fully resolve the temporal and synaptic mechanisms of its control, determine the relevant cell types involved, and define how cortical, hippocampal, and amygdalar inputs precisely influence PL activity during decision-making^13,17,21,27,59^. Dissecting whether this pathway acts via excitation, inhibition, or disinhibition will refine our understanding of how learned threats shape goal-directed actions. Moreover, one limitation of our study is the exclusive use of male rats^60^; future studies should address potential sex differences in corticostriatal circuits involved in adaptive restraint during motivational conflict. Finally, while our semi-naturalistic task provides ecological validity, extending analyses to humans through neuroimaging will be critical for assessing the translational relevance of the corticostriatal dynamics described here and for guiding strategies to modulate maladaptive decision-making. Together, these considerations place our findings within a broader framework for understanding adaptive control under motivational conflict.

## Methods

### Animals

Adult male Wistar rats (280–300 g; breeding colony at the Institute of Cellular Physiology, UNAM) were housed individually under a 12 h light/dark cycle, handled daily, and provided ad libitum water. Rats were food-restricted (12 h/day) on a controlled chow diet with a 5 g weekly bonus to maintain stable motivation for 45 mg sucrose pellets (Bio-Serv). All experiments were performed during the light phase (08:00–18:00) in compliance with the National Ministry of Health guidelines and approved by the UNAM Institutional Animal Care and Use Committee. Experiments were conducted in male rats to maintain consistency with prior work and minimize variability related to sex-dependent stress responses.

#### Behavioral tasks

All paradigms were adapted from published protocols^17,24^, and are summarized below.

### Crossing-Mediated Conflict (CMC) task

The task was conducted in two stainless-steel straight alleys (100 × 30 × 50 cm) inside a sound-attenuating chamber (150 × 70 × 140 cm). Each alley had two “safe” zones (20 × 30 cm; acrylic floor, lever, cue light, speaker, pellet dispenser, food plate) and a central “danger” zone (60 × 30 cm; grid floor delivering scrambled shocks; Coulbourn Instruments). Events (pellet delivery, cues, shocks, and lever presses) were controlled using custom MATLAB scripts. Floors and walls were cleaned between rats with soapy water and 70% ethanol. Rats were first trained to press a lever for 45 mg sucrose pellets (Bio-Serv) delivered upon cue light presentation. Each session began and ended with a 5-min context-only exposure (no cues or shocks). Conflict training lasted 34 days and comprised three phases: reward, threat, and discrimination.

#### Reward training

Rats confined to a safe zone learned to associate a light cue with food delivery. Lever presses during light-on (cue) trials delivered a pellet, whereas presses during light-off (no-cue) trials had no effect on pellet delivery. Once the response was stable, the rats were trained to cross the alley from one safe zone to an other in response to the light cue. Trials ended when the rat crossed and pressed the lever or after 180 s had elapsed. One to three lever presses were reinforced per trial to prevent habitual, non-signaled crossings.

#### Threat training

While confined to the danger zone, rats received five pairings of white noise (85 dB, 30 s) co-terminating with a scrambled foot shock (0.5 mA, 1 s) with variable intertrial intervals (1–3 min). The following day, two noise-alone trials were conducted to test threat memory. In subsequent sessions, rats were required to cross the alley for a light-cued reward, but now under threat: noise-signaled foot shock delivery (0.5 mA) in the danger zone during conflict trials.

#### Discrimination training

Short acrylic barriers (9 cm tall) were placed between the safe zones and the grid to define a “choice” zone (Fig.1A). Each session included three blocks of 10 trials, mixing non-conflict (light cue without threat) and conflict (light cue + threat) trials. The rats were allowed 180 s to decide whether or not to cross. The shock intensity and conflict trial probability progressively increased to ensure robust threat memory and maintain motivational conflict. After surgery and recovery, the rats underwent four more sessions, with seven non-conflict and three conflict trials per block, reaching 0.8 mA shock intensity by the last day. Rats that failed to discriminate between trial types (P > 0.05) or failed to cross within 180 s were excluded from the study.

#### Conflict test

The conflict test was conducted on the day after the final discrimination session. Rats were exposed to 10 crossing trials under the same conditions as the last discrimination session: seven non-conflict trials and three conflict trials (30% probability) presented in random order, with no foot shocks delivered. For optogenetic experiments, the rats completed two consecutive 10-trial blocks (nine non-conflicts and one conflict per block). In the photoinhibition sessions, the laser was activated during one non-conflict and one conflict trial in the first block, starting at trial onset and lasting until the rat crossed (∼90 s) or until 180 s had elapsed. The second block was performed without laser stimulation. In the photoactivation sessions, the first block was unstimulated, and laser stimulation was delivered during the second block following the same protocol. For single-unit recordings, the rats completed six (± 2) consecutive 10-trial blocks (nine non-conflict and one conflict trial each), all without shock delivery. Behavioral performance and neural activity were compared across conflict and non-conflict trials by analyzing the temporal and spatial dynamics of crossing, cue processing, reward-seeking, and threat responses.

### Foraging-Mediated Innate Conflict Task

This task was conducted in a custom open-field arena (90 × 90× 60 cm) placed in a dark room. The arena was divided into a peripheral “safe” zone and a central “threat” zone, brightly illuminated by an overhead light (1,500 lx). This illumination enhances natural aversion to open spaces without affecting the periphery. The arena was cleaned using soapy water and 70% ethanol between rats.

#### Conflict test

separate groups of rats underwent either a conflict or non-conflict version of the task. In the conflict condition, 30 sucrose pellets (1.35 g total) were placed in the illuminated center (“threat zone”) to introduce a motivational conflict between innate avoidance and food-seeking behavior. In the non-conflict condition, no food was placed in the center. To sustain food motivation, 10 sucrose pellets (45 mg each; 0.45 g total) were placed in each of the four corners of the arena. To reduce neophobia, rats were familiarized with sucrose pellets in their home cages (20 pellets per day for two days) before testing. For optogenetic experiments, the laser was alternated ON and OFF throughout the session in 2-minute epochs. In all experiments, each rat was individually placed in the “safe” zone and allowed to explore for 10 min. The time spent in the threat and safety zones was recorded.

### Threat Memory Retrieval Test

Rats were trained and tested in standard conditioning chambers (Coulbourn Instruments, USA). On day 1, they received five auditory fear-conditioning trials in which a tone (75 dB, 30 s) was co-terminated with a footshock (0.7 mA, 1 s) delivered through a stainless-steel grid floor. On day 2, threat memory was evaluated during two-tone presentations without shock. Each session began with a 5-min acclimation period, followed by tone trials separated by a variable inter-trial interval of 120 s. In optogenetic experiments, the laser was activated at tone onset during the first trial and remained on for a 30-s tone duration; the second tone was presented without laser stimulation. Chambers were cleaned between the rats with soapy water and 70% ethanol.

### Reward Memory Retrieval Test

Rats were placed in the safe zone of the CMC apparatus and trained for four consecutive days to press a lever for food pellets (45 mg, dustless precision pellets, Bio-Serv) on a fixed ratio 1 (FR1) schedule. On day 5, reward memory was assessed under the same conditions. In optogenetic experiments, the laser was activated during the first minute of the session and turned off during the second minute. The apparatus was cleaned with soapy water and 70% ethanol between animals.

### Low Conflict Retrieval Test

Rats were trained and tested in standard conditioning chambers (Coulbourn Instruments, USA). During the initial sessions, they learned to press a lever for sucrose pellets (45 mg, dustless precision pellets, Bio-Serv) on a variable interval 60-s (VI-60) schedule. After stable performance, they underwent aversive conditioning with five tone–shock pairings (75 dB tone, 30 s; co-terminating with a 0.7 mA, 1 s footshock). The following day, a low-conflict test was conducted with two tones presented without shock while lever pressing remained available. Each session began with a 5-min baseline and included a variable inter-trial interval of 120 seconds. In the optogenetic experiments, the laser was activated during the first tone (30 s) and remained off during the second tone. Chambers were cleaned between the rats with soapy water and 70% ethanol. During this test, rewards were immediately accessible via lever pressing, and no threat-zone navigation was required. Accordingly, this paradigm was operationally defined as a low-conflict condition relative to the CMC task, allowing assessment of threat-induced suppression of instrumental behavior independent of spatial navigation demands.

### Free Food Intake Test

To assess unconditioned feeding, rats were presented in their home cages with 10 g of sucrose pellets (45 mg, dustless precision pellets, Bio-Serv) for 15 min. For optogenetic experiments, 5 g were provided for 4 min, with laser delivery alternated ON and OFF every minute (ON during the first and third minutes; OFF during the second and fourth minutes). The pellet weight was measured before and after the session to calculate the total intake.

### Open Field Test

Rats were individually placed in a dimly lit open-field arena (20 lx) for 10 min. Locomotor activity was quantified as the total distance traveled, and anxiety-like behavior as the percentage of time spent in the central area. For optogenetic experiments, the laser was alternated ON and OFF in 2-min epochs. The arena was cleaned with soapy water and 70% ethanol between sessions.

### Stereotaxic surgery

Rats were anesthetized with isoflurane (2–3% in oxygen) and placed in a stereotaxic frame (Kopf Instruments). For pharmacological inactivation, stainless-steel guide cannulae (23-gauge) were bilaterally implanted targeting the prelimbic cortex (PL: +3.0 mm AP, ±0.6 mm ML, −2.5 mm DV), infralimbic cortex (IL: +2.8 mm AP, ±3.1 mm ML, −3.8 mm DV; 30° angle), anterior nucleus accumbens core (aNAcC: +2.5 mm AP, ±1.6 mm ML, −5.3 mm DV), or posterior NAcC (pNAcC: +1.6 mm AP, ±2.0 mm ML, −6.0 mm DV) relative to bregma, with tips positioned 1.0 mm above target. For optogenetics, AAV vectors were infused bilaterally into the PL (+3.2 mm AP, ±0.6 mm ML, −2.5 mm DV from dura) using a glass micropipette connected to a Nanoject-II injector (Drummond; 0.5 µL/site at 110 nL/min) followed by a 10-min diffusion period before retraction; optical fibers (200 µm core, 0.22 NA; Doric Lenses) were implanted bilaterally over aNAcC (+2.0 mm AP, ±3.5 mm ML, −6.4 mm DV; 16° angle) and secured with dental acrylic anchored to skull screws. For single-unit recordings, a custom movable microdrive with a 16-channel array of 35 μm tungsten microwires (California Fine Wire Co.) was unilaterally implanted in the PL; in a subset for optrode validation, Arch-expressing PL neurons were targeted with a fixed 16-wire optrode into the aNAcC (+2.5 mm AP, ±1.6 mm ML, −5.3 mm DV). All rats recovered for at least seven days before behavioral testing or recording.

### Local drug infusion

To transiently and reversibly inactivate the target regions, a cocktail of GABA_A_ and GABA_B_ receptor agonists, muscimol and baclofen (Sigma-Aldrich), was freshly prepared in sterile physiological saline on the day of infusion. Each rat received 0.5 μL per side (250 ng of each compound) delivered 15 min before behavioral testing at a rate of 0.4 μL/min using a programmable microinfusion pump (KD Scientific) connected via polyethylene tubing to a 10 μL Hamilton syringe; injectors were left in place for 2 min post-infusion to facilitate diffusion. Control rats received equivalent saline vehicle infusions, following the same procedure.

### Viral vectors

Viral vectors were obtained from the University of North Carolina Vector Core or Addgene and included AAV5-CaMKII-eArch3.0-eYFP (UNC Vector Core; titer: 3.4 × 10¹² particles/mL), AAV5-CaMKII-eYFP (UNC Vector Core; titer: 3.6 × 10¹² particles/mL), and AAV5-CaMKIIa-hChR2(H134R)-eYFP (Addgene; titer: 2.2 × 10¹³ particles/mL). All vectors were stored at −80 °C and used undiluted for microinjections.

### Laser delivery

For optogenetic manipulations, rats expressing the light-driven proton pump archaerhodopsin-3.0 (Arch 3.0) or channelrhodopsin-2 (ChR2) in the PL neuronal bodies received bilateral laser stimulation in the aNAcC. Green diode-pumped solid-state (DPSS) lasers (532 nm, continuous light, 11 mW at fiber tips, adjusted as needed within a 10–12 mW range) were used for Arch-mediated photoinhibition, whereas blue DPSS lasers (473 nm, 15 Hz, 5 ms pulse width, 12 mW, adjusted as needed within a 10–14 mW range) were used for ChR2-mediated photoactivation, following previously described protocols^23^. Laser delivery was time-locked to specific behavioral events (see Behavioral Tasks). Light was transmitted through a laser coupler/shutter (FC/PC; OEM Laser Systems), patch cords (62.5 μm core; Precision Fiber Products), a rotary joint (Doric Lenses), a dual patch cord, and custom-made bilateral optical fibers (assembled in-house from Thorlabs and Precision Fiber Products components). Rats were habituated to the tethering apparatus for three days before the first laser session to minimize stress and movement artifacts.

### Multichannel unit recordings and unit analysis

Extracellular recordings from freely behaving rats performing the conflict test were obtained using a custom-built 16-channel microdrive and Omniplex acquisition system (Plexon). Signals exceeding the voltage threshold were amplified (×100), digitized at 40 kHz, and stored for offline analysis. Spike sorting was performed with principal component analysis and template matching in an Offline Sorter (Plexon) using a semi-automated pipeline: an initial valley-seeking scan algorithm per channel followed by manual verification. Units were classified as single neurons if they formed well-isolated clusters in the principal component space and exhibited a refractory period of >1 ms; only units stable throughout the session were included. Spike timestamps and behavioral markers were exported to NeuroExplorer (NEX Technologies) for further analysis. Peri-stimulus time histograms (PSTHs; 300 ms bins) were aligned to the crossing event, defined as the moment the rat’s entire body moved from the choice to the danger zone, and averaged across trials. Firing rates were z-scored in 300 ms bins relative to a −20 to −1 s baseline, chosen to capture potential ramping before decisions. Task-modulated neurons were identified by comparing peri-event (−20 to +20 s) z-scores to baseline using two-tailed Wilcoxon signed-rank tests and classified as conflict-responsive, non-conflict-responsive, responsive in both, or unresponsive. Category distributions were compared using chi-squared tests and pairwise contrasts. Decision-period activity was assessed in a ±600 ms window around crossing, classifying neurons as excitatory if z > 2.58 (P < 0.01, two-tailed) or inhibitory if z < −1.96 (P < 0.05, two-tailed). These criteria refined the inclusion criteria for peri-event comparisons and polarity assessments. Population-level comparisons between conflict- and non-conflict-responsive neurons used two-tailed unpaired t-tests on mean z-scores in the ±600 ms window. The response magnitude was quantified as the area under the curve (AUC) for the same window, and proportions of significantly excited or inhibited neurons in each condition were derived from AUC-based classifications.

### Optrode recordings

To validate the optogenetic modulation of PL terminals in the aNAcC, rats expressing Arch in the PL were implanted with optrode arrays in the aNAcC four weeks after viral delivery. Each optrode consisted of 16 tungsten microwires (35 µm diameter; California Fine Wire Company) arranged around a central optical fiber (200 µm core, 0.22 NA). Single units were recorded while applying a repeated laser-pulse protocol of 10 alternating 20-s epochs: 10 s of continuous green laser photoinhibition (532 nm, ∼10–12 mW at the fiber tip) followed by 10 s without light. Spiking data were acquired and stored for offline spike sorting (Offline Sorter, Plexon) and spike train analysis (NeuroExplorer, NEX Technologies). Firing rate changes were assessed by comparing spike counts during laser ON versus OFF epochs using Wilcoxon signed-rank tests (1 s bins). For each 1 s bin, z-scores were calculated relative to baseline activity during the preceding 10 s OFF period, with units classified as excited if z > 2.58 (P < 0.01) or inhibited if z < –1.96 (P < 0.05) during laser activation.

### Histology

At the conclusion of behavioral experiments, rats were deeply anesthetized with sodium pentobarbital (150 mg/kg, i.p.) and transcardially perfused with 0.9% saline followed by 4% paraformaldehyde (PFA) in phosphate-buffered saline (PBS). Brains were extracted, post-fixed overnight in 4% PFA at 4 °C, and cryoprotected in 30% sucrose in PBS until saturated. Coronal sections (50 µm) were sliced using a cryostat (Leica CM1520) and processed for Nissl staining. Sections were examined under a bright-field microscope (Nikon H550S) to verify the cannula tip locations, electrode array placements, and viral expression sites. Only animals with confirmed placements within the intended target structures were included in the analyses.

### Immunohistochemistry

Rats were perfused with 0.9% saline followed by 4% paraformaldehyde (PFA) 90 min after the conflict or non-conflict test sessions. Brains were post-fixed overnight in PFA at 4 °C and then cryoprotected in 30% sucrose in PBS. Coronal sections (40 µm) were sliced using a cryostat (Leica CM1520) at multiple rostrocaudal levels of the medial prefrontal cortex and nucleus accumbens. For c-Fos immunostaining, the sections were rinsed six times in TBS with 0.1% Tween-20, incubated in 3% hydrogen peroxide (Sigma) for 10 min to quench endogenous peroxidase activity, and blocked for 2 h at room temperature in 4% normal goat serum (Vector Laboratories), 4% bovine serum albumin (Santa Cruz Biotechnology), and 0.1% Triton X-100 in TBS-Tween. Sections were then incubated for 48 h at room temperature with rabbit polyclonal anti-c-Fos (1:1000; Millipore), followed by a 1 h incubation with biotinylated goat anti-rabbit secondary antibody (1:500; Jackson ImmunoResearch). The s ignal was amplified using the ABC Elite Kit (Vector Laboratories) and visualized using DAB (SK-4100; Vector Laboratories), after which the tissues were washed three times in PBS. For NeuN immunostaining, the same protocol was applied using rabbit anti-NeuN conjugated with Alexa Fluor 647 (1:2000; Abcam). Sections were mounted on gelatin-coated slides and coverslipped with aqueous mounting medium.

### Microscopy and image analysis

For bright-field microscopy, c-Fos-labelled neurons were quantified automatically using ImageJ (RRID: SCR_003070). Sections were imaged at 10× magnification with a Nikon H550S microscope equipped with a digital camera, acquiring micrographs at three anterior–posterior levels from bregma for each region of interest: medial prefrontal cortex (+3.5 to +2.5 mm) and nucleus accumbens (+2.5 to +1.5 mm). Neurons were classified as c-Fos positive if the nucleus was clearly distinguishable from the background, with an area of 25–250 μm² and circularity ≥ 0.60. A consistent, predetermined intensity threshold was applied across all samples using ImageJ’s automated particle analysis function to ensure objective identification of positive nuclei. For each region, the labeled cells were counted in three sections per rat from one hemisphere and averaged. For optogenetic experiments, fluorescence images of NeuN-immunostained tissues were acquired using a confocal/two-photon microscope (Zeiss LSM710) at 10×, 20×, and 40× magnification. Z-stacks of 18 optical sections were collected at 3 μm intervals, spanning ∼50 μm of tissue per region; maximum intensity projections were generated along the Z-axis, and channels were merged. Representative images of viral expression were obtained using TILE scans at 10× or 20× magnification, with a pinhole of ∼10 μm. All images were processed using ImageJ, pseudo-colored according to the fluorophore, and adjusted for brightness and contrast.

### Data collection and analysis

All behaviors were recorded using overhead digital video cameras (Provision D-380D5) positioned above each apparatus and analyzed using ANY-maze 7.1, an automated tracking software (Stoelting, USA). The straight alley was divided into four zones (start, 19 cm; choice, 10 cm; danger, 42 cm; goal, 29 cm) to assess temporal and spatial behavioral patterns, including cue responses, crossings, reward-seeking, and defensive reactions. Tracking at three body points (head, center, tail) provided data on time spent and entries per zone; the center point determined presence in the start, danger, and goal zones, whereas the head point determined the choice zone presence (within 10 mm for a successful visit). The percentage time per zone was calculated as: (time per zone × 100) / total time in the start, danger, and goal zones. Additional measures included freezing bouts (≥300 ms immobility in the start zone), latency to press (lever on the same side), risk assessment events (choice-zone entries with approach/avoidance oscillations toward the reward site), crossing speed (m/s across danger zone), and latency to press in the goal zone (sum of times in start, danger, and goal zones). Movement heatmaps were generated from head tracking with a 10-s maximum for the hottest color. In all tests, freezing was defined as ≥300 ms immobility (excluding respiration) during auditory cues and was expressed as a percentage of cue duration. Distance traveled and center time in the open field were automatically computed in the ANY-maze. Food intake was calculated as follows: (food consumed × 100) / food offered, with pellet weights measured (each pellet weighed 45 mg) before and 30 min after access; intake was manually recorded by experimenters blinded to the condition. Data were analyzed using unpaired/paired two-tailed t-tests and mixed-design two-way ANOVAs, followed by Bonferroni post hoc tests. Planned comparisons within trial types were conducted regardless of interaction, based on predefined hypotheses. Analyses were performed using STATISTICA (StatSoft, USA) and GraphPad Prism 7, with significance set at P < 0.05.

## Acknowledgments

We thank Dr. Christian Bravo-Rivera for his comments on the previous version of the manuscript and Dr. Leticia Ramírez-Lugo for her technical assistance. Erick López Roldán for assistance in the design of the microdrive, and Sotres-Bayon laboratory members for helpful discussions.

## Author Contributions

Conceptualization: E.I-H, F. S-B

Methodology: E.I-H, F.S-B.

Experimental work: E.I-H, E.H-O.

Data analysis: E. I-H, F. S-B.

Resources: F. S-B

Supervision: F. S.-B.

Writing: E.I-H. and F. S.-B.

## Data and materials availability

All data needed to evaluate the conclusions in the paper are present in the main text and/or the Supplementary Materials. Additional data related to this paper may be requested from the authors.

