## supplementary materials for "Top-down corticostriatal gating of adaptive restraint during motivational conflict"

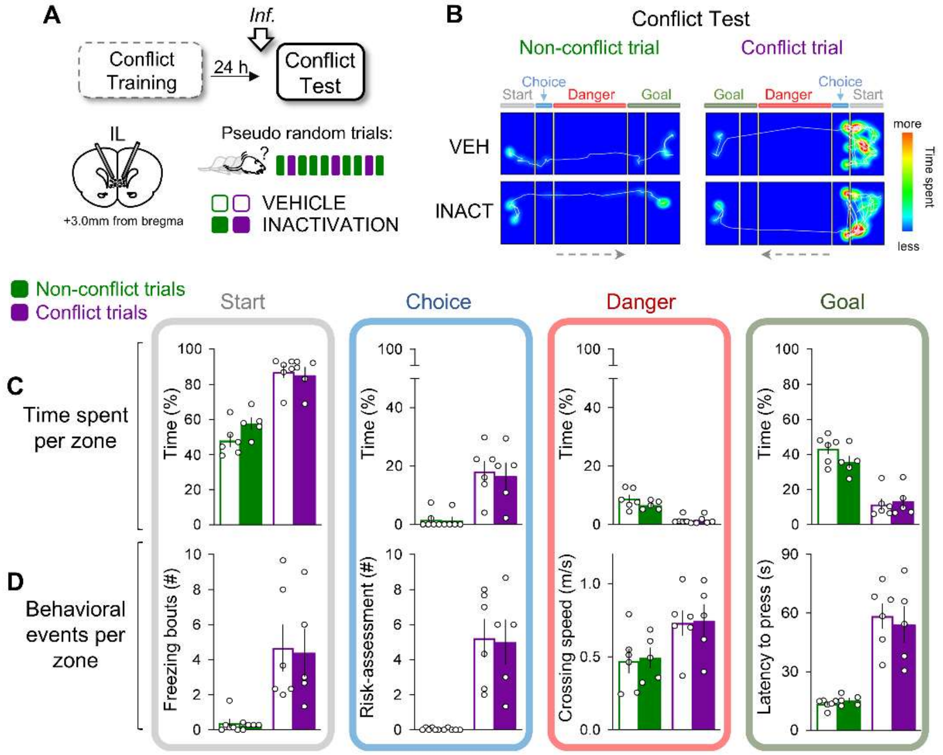


**Fig. S1. IL inactivation does not affect restraint under learned threat.**

(**A**) Bilateral cannula placements in the infralimbic cortex (IL). Rats received vehicle (VEH, n = 6) or muscimol + baclofen (INACT, n = 5) before the conflict test.

(**B**) Representative tracks and heat maps during conflict (reward + threat) and non-conflict (reward-only) trials.

(**C**) IL inactivation did not significantly change time in the start ($F(1,9) = 1.31$, $P = 0.28$), choice ($F(1,9) = 0.09$, $P = 0.76$), danger ($F(1,9) = 0.96$, $P = 0.35$), or goal zones ($F(1,9) = 1.04$, $P = 0.33$) across trial types.

(**D**) Defensive behaviors—freezing ($F(1,9) = 0.04$, $P = 0.84$), risk assessment ($F(1,9) = 0.02$, $P = 0.88$), velocity ($F(1,9) = 0.04$, $P = 0.84$), and press latency ($F(1,9) = 0.05$, $P = 0.82$) were unaffected by IL inactivation in both trial types.

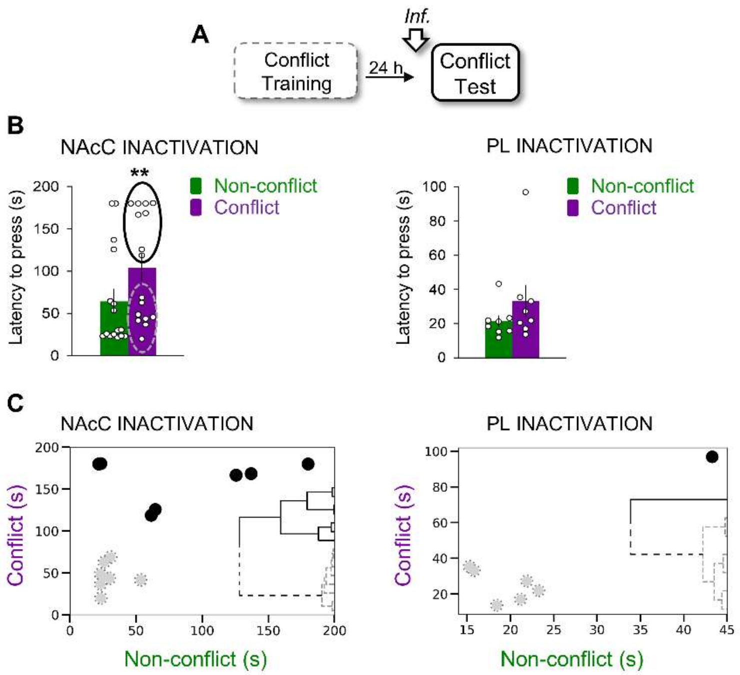


**Fig. S2. The anterior–posterior organization of the NAcC underlies the divergent effects of inactivation on behavior.**

(**A**) Experimental timeline for NAcC (n = 16) and PL (n = 8) inactivation before the conflict test.

(**B**) NAcC inactivation produced heterogeneous behavioral effects, with significant differences between conflict and non-conflict latencies ($t(15) = 3.15$, $P = 0.006$), unlike the uniform pattern observed after PL inactivation ($t(7) = 1.74$, $P = 0.12$).

(**C**) Unsupervised clustering revealed two NAcC subgroups (gray vs. black) corresponding to anterior (+2.5 to +2.0 mm) and posterior (+2.0 to +1.6 mm) infusion sites. PL inactivation did not show any clear subgroups.

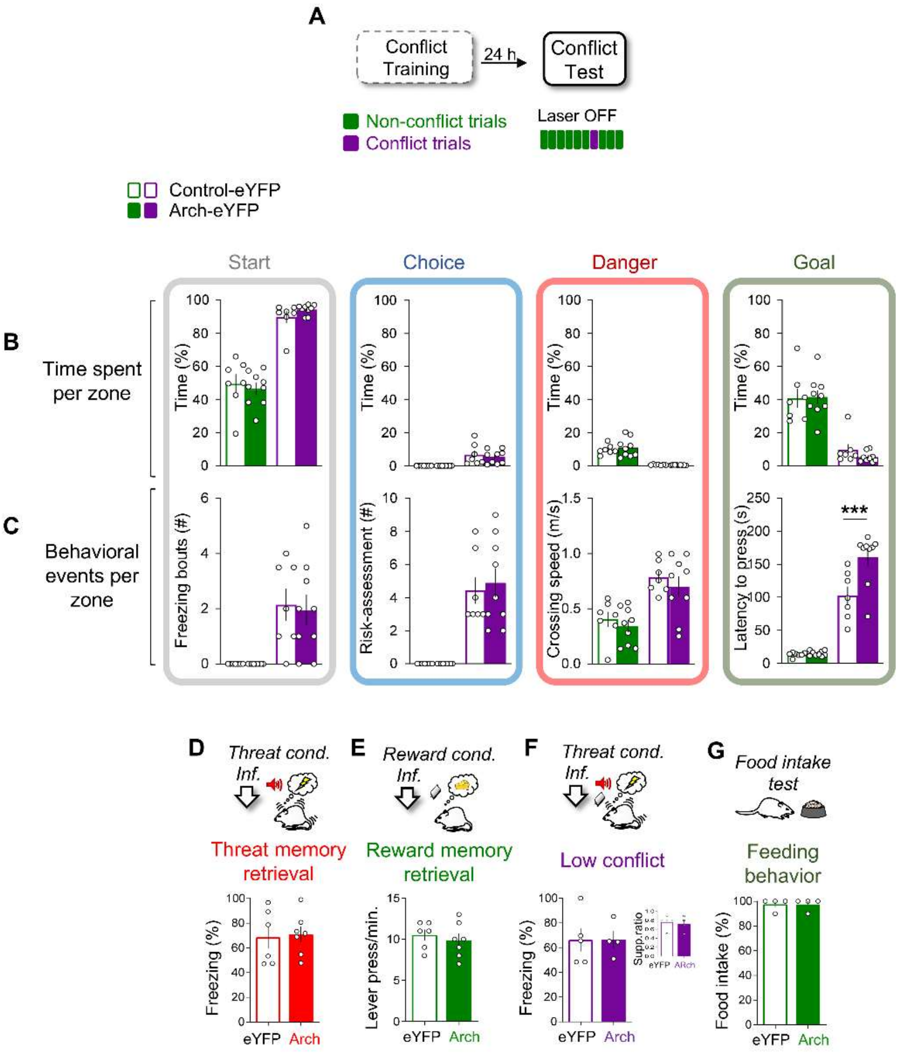


**Fig. S3. Conflict test trials without optogenetic stimulation (laser off).**

(**A**) After the laser ON block, rats completed a subsequent block of laser-OFF trials during the conflict test (Control-eYFP, n = 7; Arch-eYFP, n = 9).

(**B**) During laser-OFF trials, Arch-expressing rats spent similar times in the start ($F(1,14) = 0.06$, $P = 0.80$), choice ($F(1,14) = 0.26$, $P = 0.61$), danger ($F(1,14) = 0.18$, $P = 0.67$), and goal zones ($F(1,14) = 0.24$, $P = 0.63$) as controls.

(**C**) Arch rats showed longer crossing latencies ($F(1,14) = 9.30$, $P = 0.008$; conflict trials $P = 0.0004$) but no differences in freezing ($F(1,14) = 0.05$, $P = 0.81$), risk assessment ($F(1,14) = 0.13$, $P = 0.71$), or crossing speed ($F(1,14) = 0.95$, $P = 0.34$).

(**D–E**) No group differences were found in freezing ($t(11) = 0.19$, $P = 0.84$) or lever pressing ($t(11) = 0.60$, $P = 0.55$) during threat (Control-eYFP, n = 6; Arch-eYFP, n = 7) or reward (Control-eYFP, n = 6; Arch-eYFP, n = 7) memory retrieval tests.

(**F**) In the low-conflict test without stimulation (Control-eYFP, n = 5; Arch-eYFP, n = 4), freezing ($t(7) = 0.009$, $P = 0.99$) and suppression ratios ($t(7) = 0.32$, $P = 0.75$) were similar between groups.

(**G**) In the free food intake test without stimulation (Control-eYFP, n = 4; Arch-eYFP, n = 4), food intake was comparable ($t(6) = 0.00$, $P > 0.99$). Data are mean ± SEM; ***$P < 0.001$, Bonferroni post hoc test following two-way mixed-design ANOVA or Student's t-test.

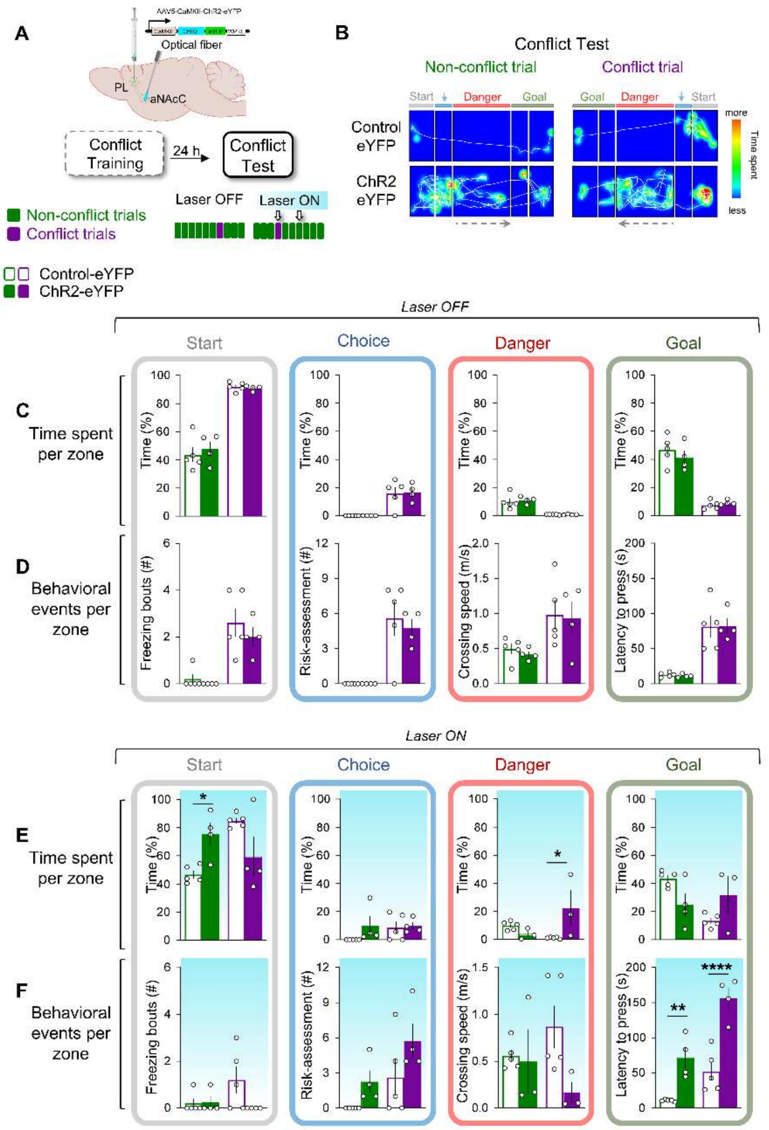


**Fig. S4. Activating PL→aNAcC projection disrupts goal-directed behavior regardless of threat.**

**(A)** Schematic representation of AAV5-CaMKII-ChR2-EYFP injection in the PL and bilateral optical fiber implantation in the aNAcC. During the conflict test (Control-eYFP, n = 5; ChR2-eYFP, n = 4), rats completed two trial blocks: laser OFF followed by laser ON.

(**B**) Representative movement tracking of eYFP and ChR2 rats during conflict and non-conflict trials.

(**C**) In laser-OFF trials, ChR2 rats spent a similar time in the start ($F(1,7) = 0.16$, $P = 0.70$), choice ($F(1,7) = 0.01$, $P = 0.89$), danger ($F(1,7) = 0.15$, $P = 0.70$), and goal ($F(1,7) = 0.24$, $P = 0.63$) zones compared to controls.

(**D**) No differences were observed in freezing ($F(1,7) = 1.31$, $P = 0.32$), risk assessment ($F(1,7) = 0.21$, $P = 0.65$), crossing speed ($F(1,7) = 0.16$, $P = 0.69$), or press latency ($F(1,7) = 0.001$, $P = 0.97$).
(**E**) Photoactivation produced no main effect of group but a significant interaction, increasing start-zone occupancy during non-conflict trials (interaction: $F(1,7) = 12.08$, $P = 0.01$; $P = 0.03$) and danger-zone occupancy during conflict trials (interaction: $F(1,6) = 10.69$, $P = 0.01$; $P = 0.018$).
(**F**) Photoactivation significantly prolonged press latency in both trial types ($F(1,7) = 32.97$, $P = 0.0007$; conflict $P < 0.0001$, non-conflict $P = 0.005$) but did not affect freezing ($F(1,7) = 2.06$, $P = 0.19$), risk assessment ($F(1,7) = 4.00$, $P = 0.08$), or crossing speed ($F(1,6) = 4.46$, $P = 0.07$). Data are mean ± SEM; *$P < 0.05$, **$P < 0.01$, ****$P < 0.0001$ in Bonferroni post hoc tests following two-way mixed-design ANOVA.

**Movie S1. Representative behavior during conflict and non-conflict trials in the conflict test.**

Automated video tracking of a rat performing the crossing-mediated conflict task during the conflict test (no shocks delivered), shown at 2× speed. The video depicts successive trials in which the animal decides whether to cross an electrified grid to obtain food. In conflict trials, the reward cue (reward light ON) is presented together with the danger cue (white noise LED ON), requiring the animal to withhold approach in the face of potential threat. In non-conflict trials, only the reward cue is presented, allowing safe reward pursuit. Trajectories and zone occupancy were recorded using automated tracking software.
